# TLR-induced reorganization of the IgM-BCR complex regulates B-1 cell responses to infections

**DOI:** 10.1101/595694

**Authors:** Hannah P. Savage, Kathrin Kläsener, Fauna L. Smith, Zheng Luo, Michael Reth, Nicole Baumgarth

**Affiliations:** Center for Comparative Medicine, University of California, Davis, Davis, CA 95616, USA; Graduate Group in Immunology, University of California, Davis, Davis, CA 95616, USA; Integrated Pathobiology Graduate Group, University of California, Davis, Davis, CA 95616, USA; BIOSS Centre for Biological Signalling Studies, University of Freiburg, D-79104 Freiburg, Germany; Department of Molecular Immunology, Institute of Biology III at the Faculty of Biology of the University of Freiburg, D-79104, and at the Max Planck Institute of Immunobiology and Epigenetics, D-79108 Freiburg, Germany; Dept. Pathology, Microbiology and Immunology, School of Veterinary Medicine, University of California, Davis, CA 95616, USA

## Abstract

Neonatally-developing, self-reactive B-1 cells generate steady levels natural antibodies throughout life. They can, however, also rapidly respond to infections with increased local antibody production. The mechanisms regulating these two seemingly very distinct functions are poorly understood, but have been linked to expression of CD5, an inhibitor of BCR-signaling. Here we demonstrate that TLR-mediated activation of CD5+ B-1 cells induced the rapid reorganization of the IgM-BCR complex, leading to the eventual loss of CD5 expression, and a concomitant increase in BCR-downstream signaling, both *in vitro* and *in vivo* after infections with influenza virus and *Salmonella typhimurium*. Both, initial CD5 expression and TLR-mediated stimulation, were required for the differentiation of B-1 cells to IgM-producing plasmablasts after infections. Thus, TLR-mediated signals support participation of B-1 cells in immune defense via BCR-complex reorganization.

## Introduction

During lymphocyte development (self)-antigen binding by the TCR and BCR results in negative selection, leading to the removal of strongly self-reactive lymphocytes from the T and B cell repertoire. Depending on the strengths of these antigen-BCR interactions, self-reactive B cells undergo deletion, receptor-editing, or they become anergic, i.e. unresponsive to antigen-receptor engagement ^1^.

Self-reactive, anergic bone marrow-derived B cells up-regulate expression of the signaling inhibitor CD5 ^2^. On developing T cells, the levels of CD5 expression correlate with TCR signaling intensity encountered during thymic development, with those most strongly binding to self-antigens expressing the highest levels of CD5 ^3, 4^. While several ligands have been proposed for CD5 ^3, 5, 6, 7, 8, 9, 10, 11^, none seem to have significant CD5-dependent biological effects. Instead, CD5 expression seems to directly reduce antigen-receptor signaling ^2, 3, 4, 12^. Thus, CD5 seems to act primarily as a component of the antigen-receptor complex, directly modulating TCR and BCR signaling.

In addition to anergic B cells, most B-1 cells also express CD5. In contrast to lymphocytes developing postnatal, these primarily fetal and neonatal-derived B cells ^13, 14, 15^ seem to undergo a positive selection step during development, requiring self-antigen recognition and strong BCR signaling. The lack of self-antigen expression, or the deletion of co-stimulatory molecules that enhance BCR signaling, diminished B-1 cell development, while deletion of negative co-stimulatory signals, or enhanced surface expression of the BCR, resulted in enhanced development ^5, 15, 16, 17, 18^. Specificity of CD5+ B-1 cells for self-antigens and self-antigen binding during development is consistent with their known self-reactive BCR repertoire ^19, 20, 21, 22^ and thus a role for CD5 in silencing B-1 cell responses to BCR-engagement in order to avert autoimmune responses.

Yet, not all B-1 cells express CD5. Depending on their expression or not of CD5, they are typically divided into two subsets, B-1a and B-1b, respectively. Consistent with their expression of CD5, B-1a cells do not proliferate in response to BCR stimulation ^23^. However, in CD5-/- mice and in mice in which the association of membrane IgM with CD5 was inhibited, mature B-1 cells demonstrated a proliferative response similar to that of conventional B (B-2) cells ^24, 25^, further confirming that CD5 expression reduces B-1 cell responsiveness to BCR-signaling.

A BCR-signaling independent response of B-1 cells might be inferred from the fact that B-1 cells strongly respond to innate, TLR-mediated signals, such as LPS, and that they are a major source for “natural” self-reactive IgM. Moreover, steady-state secretion of natural IgM does not appear to require external antigenic stimulation, as total serum levels of natural IgM and frequencies of IgM-secreting B-1 cells are similar in mice held under both, SPF and germfree housing conditions ^26, 27^. However, natural IgM production is not stochastic, but instead likely driven by expression of self-antigens. This was demonstrated by Hayakawa et al, who showed a lack of anti-Thy-1 self-reactive IgM antibodies in the serum of Thy-1-deficient but not Thy-1 expressing mice ^19, 28^, as well as repertoire studies By Yang et al, which showed selective and extensive clonal expansion of certain CD5+ B-1 cell clones during the first 6 months of life, including in germfree mice ^22^.

Furthermore, B-1 cells can be actively involved in immune responses to various pathogens ^29, 30, 31, 32, 33^. Given that CD5 is a BCR inhibitor, it was suggested that CD5-B-1b cells, but not B-1a cells, respond to pathogen encounters in an antigen-dependent manner. Haas and colleagues, conducting studies in CD19-deficient mice that lack B-1a development, concluded that B-1a cells are responsible for natural IgM secretion, while only the B-1b cells responded to *Streptococcus pneumonia* infection ^29^. Similarly, CD5-B-1b cells were shown to expand and secrete protective IgM after infection with *Borrelia hermsii* and *Salmonella typhimurium* ^30, 31, 32^.

This model of a “division of labor” between B-1a and B-1b cells leaves the B-1 cell response to influenza infection as an outlier. Chimeric mice reconstituted with either allotypically-marked CD5+ or CD5-B-1 cells showed that only CD5+ B-1 cells were responding *in vivo* to influenza infection with migration from the pleural cavity to the draining mediastinal lymph nodes (MedLN), where they differentiated into IgM-secreting cells ^33, 34^. The reasons for the apparent different behaviors of CD5+ and CD5-B-1 cells in the various infectious disease models are unexplained. Furthermore, it is unclear how B-1 cells expressing CD5 can participate in antigen-specific immune responses.

This study addresses some of these questions and reconciles previous divergent findings on B-1 cell responses to infections by demonstrating that CD5+ B-1 cells are the responding B-1 cell population to infection with both, influenza virus as well as *Salmonella*. However, once activated, these B-1 cells lose expression of CD5 and thus become “B-1b” like. Mechanistically, downregulation of CD5 required expression of TLR, triggering of which resulted in the reorganization of the IgM-BCR complex. TLR-mediated reorganization led to a rapid dissociation, and eventual loss of CD5 from the complex, and triggered enhanced IgM-CD19 and CD79:Syk interactions, and thus activation of downstream BCR-signaling pathways. Thus, TLR-mediated signals support participation of B-1 cells in immune defense via BCR-complex reorganization, linking innate and adaptive antigen-recognition by B-1 cells.

## Results

### CD5 negative B-1 cells are responsible for local IgM secretion after influenza infection

We previously identified three populations of cells involved in natural IgM secretion: CD5+ B-1 cells, CD5-B-1 cells, and plasma cells, the latter are CD19- and CD138/Blimp-1+ ^35^. Using a neonatal chimera model, in which the B-1 cells of neonatal mice are replaced by allotype-disparate peritoneal cavity B-1 cells ^36^, we demonstrated that the natural IgM-secreting plasma cells are B-1-derived (B-1PC) ^35^. Because B-1-derived IgM is important for protection from lethal influenza infection ^37^, we sought to determine which B-1 cell populations generate IgM in the draining (mediastinal) lymph nodes (MedLN) after influenza infection ^33^.

MedLN of allotype-chimeras infected for 7 days with infection with influenza A Puerto Rico 8/34 (A/PR8) showed a rapid accumulation and then differentiation of B-1 cells to IgM-secreting B-1PC (Fig. 1A). Chimeras generated with B-1 donor cells from Blimp-1 YFP reporter mice ^38, 39^ confirmed the presence of Blimp-1-YFP+ B-1PC in the MedLN (Fig. 1B). The MedLN B-1PC mostly lacked expression of CD5, particularly among the Blimp-1hi cells (Fig. 1C). This was surprising, as we had shown previously B-1 cell migration from the pleural cavity to the MedLN and their local IgM secretion was restricted to CD5+ but not CD5-CD19+ CD43+ B-1 cells after influenza infection ^34^.

**Figure 1:**
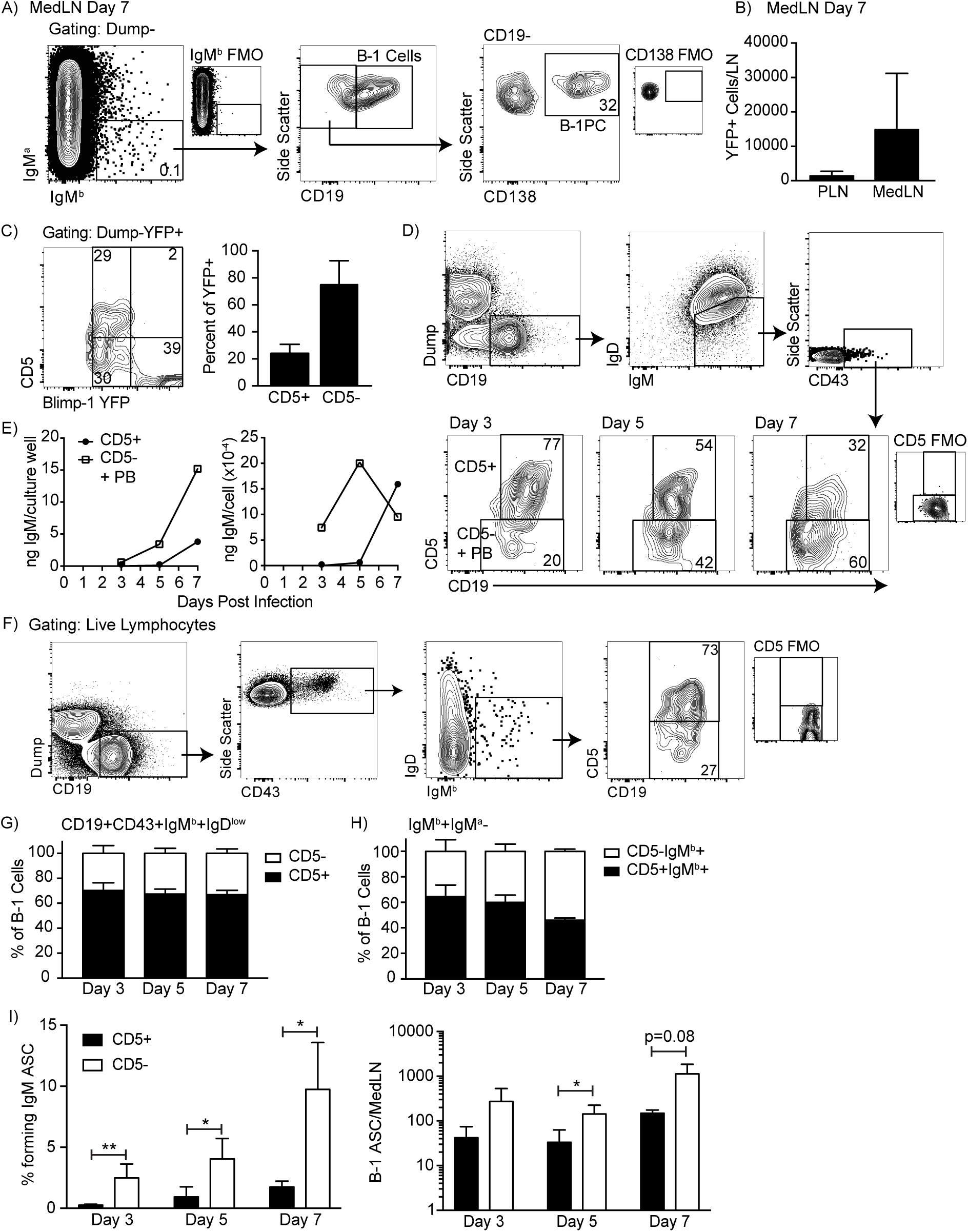
CD5 negative B-1 cells secrete most IgM in the mediastinal lymph nodes (MedLN) after influenza infection. (A) FACS plot of MedLN cells from day 7 influenza-A/PR8-infected neonatal chimeric mice generated with Ighb B-1 donor cells and Igha host cells. Shown is gating to identify IgMb+CD19+ B-1 cells and IgMb+CD19-CD138+ B-1PC. FMO, “fluorescence minus one” control stains. (B) Mean number ± SD of Blimp YFP+ cells in peripheral LN (PLN) and MedLN of day 7 influenza-infected neonatal chimera generated with B-1 donor cells from Blimp-1-YFP mice (n=4). (C) FACS plot (left) and (right) mean percentage ± SD of CD5+ and CD5-cells among total Blimp-1 YFP+ cells (n=13). (D) FACS gating strategy for sorting CD19+IgM+IgD^lo^CD43+ CD5+ or CD5-cells in the MedLN on days 3, 5, or 7 after influenza infection of C57BL/6 mice, pooled from n=2-3 per time point. (E) Concentration (ng/ml) IgM in supernatant (left) and secreted (ng × 10^−4^) per cell (right) of sorted cells measured by ELISA. (F) FACS gating strategy and (G/H) mean percentage ± SD of CD19+CD43+IgM^b^+IgD^lo^ and CD5+ or CD5-B-1 cells among (G) Dump-CD19+ IgMb+ and (H) total (Dump-IgMb+ IgMa-) B-1 cells at indicated times after infection (n=6-7 per time point). (I) Mean percentage ± SD (left) and total number ± SD (right) of FACS-sorted CD5+ and CD5-B-1 cells (IgM^b^+IgM^a^-) that formed IgM antibody-secreting cells (ASC) in each MedLN, as measured by ELISPOT (n=3-4 per time point). Results are representative of >4 (A), 3 (B), and 2 (F, I), or are combined from 2 (D, E, G, H) or 3 (C) independent experiments. Values in (I) were compared by unpaired Student’s t test (*=p<0.05, **=p<0.005).

To investigate the contribution of CD5-B cells to local IgM secretion, we FACS-sorted CD19+IgM+IgD^lo^CD43+ CD5+ and CD5-B cells on days 3, 5, and 7 after influenza infection from C57BL/6 mice (Fig. 1D), which were then cultured for 2 days to analyze spontaneous IgM secretion by ELISA. Consistent with the presence of CD5-B-1PC, cells lacking CD5 secreted more IgM compared to CD5+ cells when measuring total IgM concentrations and calculating IgM production per cell (Fig 1E). Sorted CD5+ cells did not secrete measurable amounts of IgM unless harvested after day 5 of infection.

Because CD5-B-1 cells and IgM-secreting B-2 derived plasmablasts express a similar phenotype (IgM+ IgD-CD5-CD19+ CD43+), the CD5-cells could have contained both B-1 cells and/or B-2-derived IgM-secreting cells. To determine the contribution of CD5-B-1 cells to IgM secretion in the MedLN, we infected allotype-chimeras, in which B-1 (Igh^a^) and B-2 (Igh^b^) cells and their secreted antibodies were distinguished based on Igh-allotype differences ^36^. The studies confirmed our previous findings that among CD19^hi^IgM^b^+IgD^lo^CD43+ B-1 cells in the MedLN, about 70% expressed CD5 after influenza infection (Fig. 1F-G).

Because we had shown previously that Blimp-1+ B-1PC have reduced or absent CD19-expression ^35^ and found here that these cells are present after influenza infection (Fig. 1A-B) and often lacked CD5-expression (Fig. 1C), we expanded the analysis to include all IgM^b^-expressing (B-1 donor-derived) and IgM^a^ negative (recipient-derived) cells, regardless of expression of CD19 or other surface markers. In contrast to the analysis described above, this expanded analysis of all B-1 donor Igh-b cells revealed that the frequency of CD5 negative MedLN B-1 cells increased after influenza infection (Fig. 1H), consistent with the development of CD5-B-1PC in this compartment (Fig. 1A-C). Furthermore, FACS-sorting and culture of CD5+ and CD5-B-1 cells showed that a higher frequency and total number of CD5-B-1 cells secreted IgM in the MedLN compared to CD5+ B-1 cells on days 3, 5, and 7 after infection (Fig. 1I). Thus, CD5-B-1 cells increase in the MedLN and are a major source of local IgM production after influenza infection.

### CD5+ B-1 cells decrease CD5 expression after LPS stimulation in vitro

To reconcile our previous findings about the role of CD5+ B-1 cells in influenza infection ^33, 34^, we considered whether CD5 surface expression changes after B-1 cell activation. Indeed, approximately 40% of highly purified FACS-sorted CD5+ B-1 cells from the peritoneal cavity lacked CD5 expression when cultured for 3 days in the presence but not absence of LPS, a stimuli that is known to induce IgM production by body cavity B-1 cells ^40^ (Fig. 2A). CD5 surface expression was unaffected during the first 2 cell divisions following stimulation, but was then quickly lost during the next 1-2 divisions (Fig. 2B). Both, surface-expressed CD5 and *cd5* mRNA, as assessed by qRT-PCR, were decreased among B-1 cells after 3 days of LPS stimulation (Fig. 2C-D). Surface CD5 levels were decreased first, by 1.5 days of culture, while *cd5* mRNA was not reduced until 2 days after culture onset (Fig. 2C-D). The stimulated cells began secreting IgM before CD5 levels were reduced, but the increase in IgM secretion was more pronounced after at least 2 days of stimulation compared to the earlier time points (Fig. 2E).

**Figure 2:**
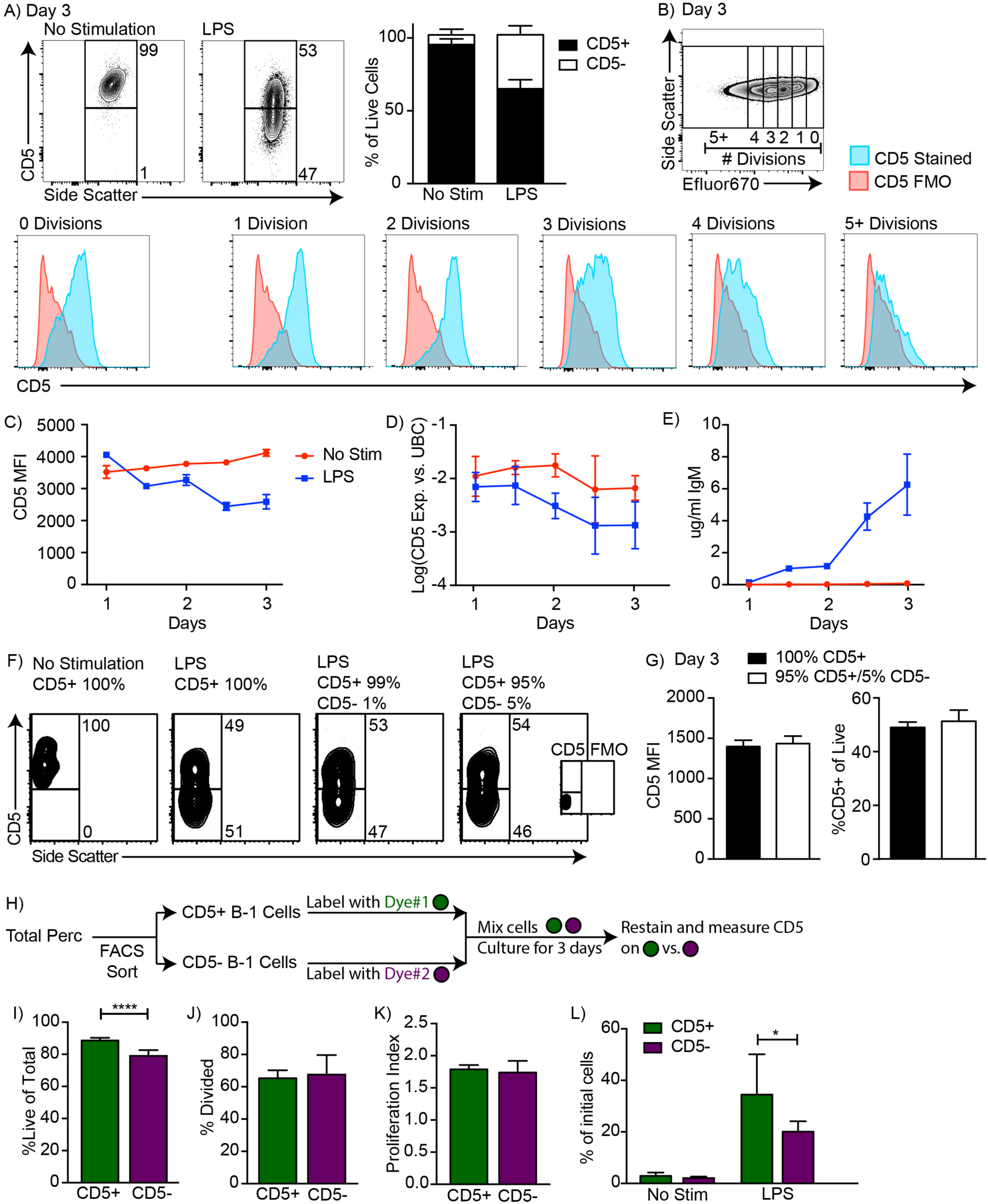
CD5+ B-1 cells decrease CD5 expression after LPS stimulation *in vitro*. (A) Representative FACS plots (left) and mean percentage ± SD (right) of CD5+ and CD5-B-1 cells after FACS-purified peritoneal cavity CD19+ CD23-CD5+ B-1 cells were cultured with or without 10 µg/ml LPS for 3 days (n=18). (B) CD5 expression on FACS-purified Efluor 670-stained proliferating peritoneal cavity CD5+ B-1 cells stimulated with LPS compared to CD5 FMO (fluorescence minus one) control. (C) Mean CD5 MFI ± SD, determined by flow cytometry, (D) mean Log(*cd5* mRNA expression) ± SD, determined by qRT-PCR, and (E) mean IgM secretion ± SD (µg/ml), determined by ELISA, after purified peritoneal cavity CD5+ B-1 cells were cultured for indicated times with LPS (n=3-4 per time and data point). (F) FACS plots from cultures with or without LPS of purified CD5+ B-1 cells mixed or not with indicated percentages of CD5-B-1 cells. (G) Mean CD5 MFI ± SD (left) and mean percentage ± SD of CD5+ cells (right) of cultures in F (n=3). (H) CD5+ (green) and CD5- (purple) B-1 cells were each labeled with either Efluor670 and CFSE. Dyes used to label each population were switched for repeated experiments. (I) Mean percentage ± SD of live cells, or (J) of divided cells, (K) mean number of divisions ± SD among cells that had divided, and (L) mean percentage ± SD of cell numbers on day 3 compared to input numbers for CD5+ and CD5-cells cultured with LPS (n=8). Results are combined from 4 (A), or are representative of >5 (B), 4 (H-L) and 2 (C - G) independent experiments, respectively. Values in (G) and (I-L) were compared using an unpaired Student’s t test (*=p<0.05, ****=p<0.00005).

A number of control studies ensured that the reduced frequencies of CD5+ B-1 cells in the cultures were due to a loss of surface expression and not to an expansion of small numbers (< 5%) CD5-cells that might have contaminated the cultures at onset. First, we separated by FACS CD5+ and CD5-B-1 cells from the body cavities to very high purities, and then cultured either 100%, 99% or 95% sorted CD5+ cells to which we added 0%, 1% and 5% CD5-cells, respectively. The percentage of CD5+ and CD5-cells after 3 days of culture was unaffected by the initial composition of the culture wells (Fig. 2F). When cultures with 100% and 95% CD5+ at onset were compared on day 3 of culture, there was no significant difference in either CD5 MFI or in the percent of CD5+ and CD5-cells (Fig. 2G). Thus, small percentages of CD5-B-1 cells at culture onset, representative of potential sort impurities, could not explain the lack of CD5 expression by the CD5+ B-1 cells stimulated with LPS for 3 days.

Next, we compared the ability of CD5+ and CD5-B-1 cells to survive and/or proliferate with and without LPS stimulation. To ensure that the two populations were exposed to the same culture conditions, CD5+ and CD5-B-1 cells were sorted, labeled with different proliferation dyes, and cultured together (Fig. 2H). Compared to B-1 cells that expressed CD5 on day 0, CD5-cells survived less well after stimulation (Fig. 2I). B-1 cells expressing or not expressing CD5 at culture onset responded similarly with proliferation to LPS stimulation in terms of the percentage of cells that underwent division as well as the numbers of divisions each B-1 cell underwent (Fig. 3J-K). Reflecting the similar rates of proliferation and the increased survival of the CD5+ B-1 cells, populations of B-1 cells that expressed CD5 at culture onset were present at higher frequencies of input cells compared to B-1 cells that were CD5 negative (Fig. 2L). Thus, we conclude that CD5+ B-1 cells lose CD5 surface and mRNA expression after *in vitro* LPS stimulation.

**Figure 3:**
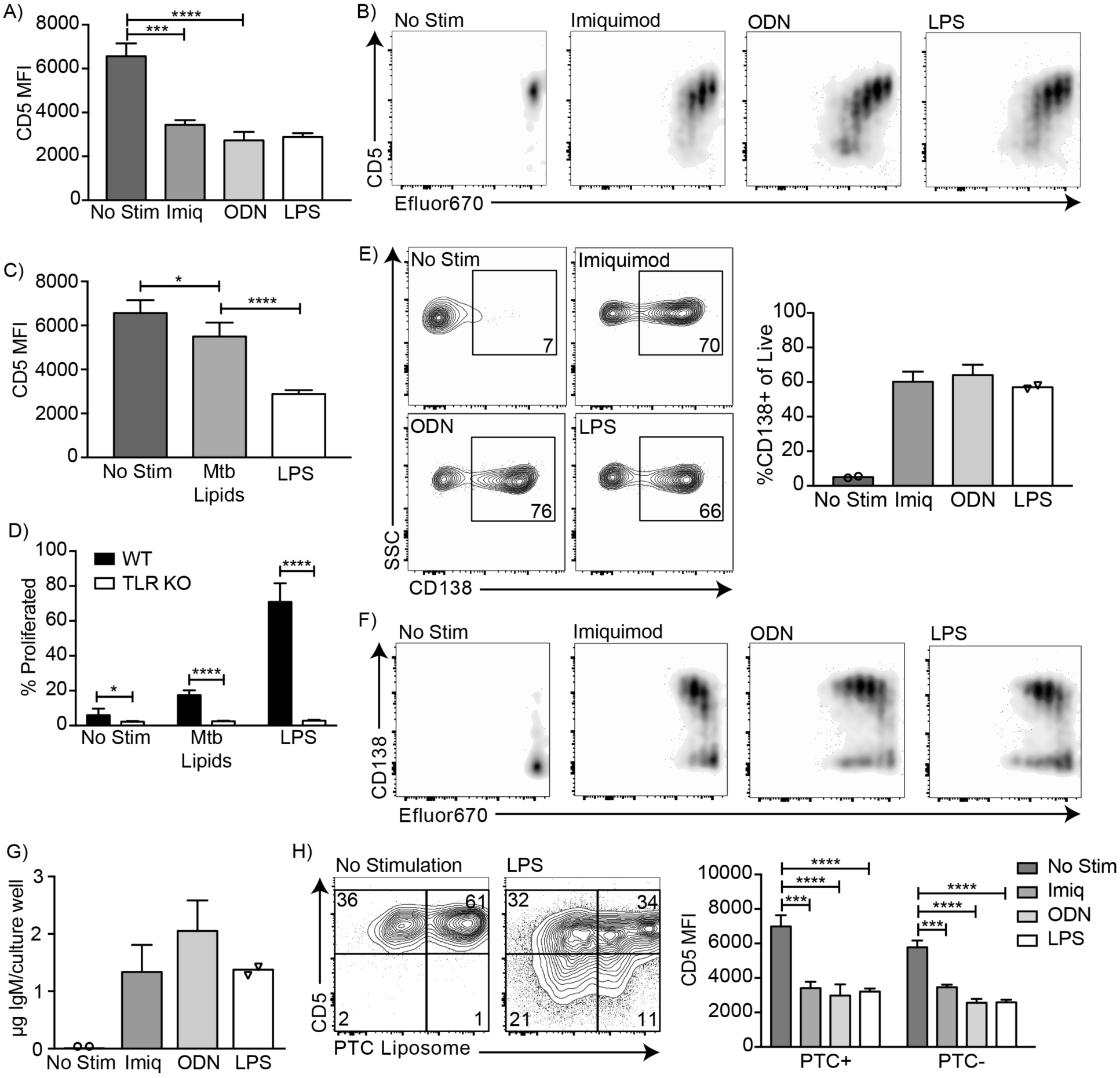
CD5+ B-1 cells differentiate into CD5-IgM secreting cells after TLR-mediated activation. (A) CD5 MFI ± SD and (B) representative FACS plots for CD5+ B-1 cells cultured without stimulation or with Imiquimod (TLR7 agonist), ODNs (CpG 7909), or LPS (n=3-5). (C) Mean CD5 MFI ± SD of CD5+ B-1 cells cultured without stimulation or with *Mycobacterium tuberculosis* (Mtb) lipids or LPS (n=4-5). (D) Mean percentage ± SD of B-1 cells from wild type (WT) or Tlr2^-/-^xTlr4^-/-^xUnc93b1^3d/3d^ (TLR KO) mice that underwent at least one division after culture without stimulation or stimulated with *Mycobacterium tuberculosis* (Mtb) lipids or LPS (n=6-9 per group). (E) FACS plots (left) and mean percentage ± SD (right) of CD138+ cells, and (F) representative FACS plots for CD138 expression among proliferating cells. (G) Mean IgM concentration ± SD (µg total per culture well) of cultured CD5+ B-1 cells stimulated or not with Imiquimod (TLR7 agonist), ODNs (CpG 7909), or LPS (n=2 for no stimulation and LPS, n=5 for Imiquimod and ODN). (H) Sample FACS plot (left) and mean CD5 MFI ± SD (right) of PTC liposome-binding (PTC+) and non-PTC liposome-binding (PTC-) cells for CD5+ B-1 cells cultured without stimulation or with Imiquimod (TLR7 agonist), ODNs (CpG 7909), or LPS (n=3-5). Results are combined from two (D, E-G), or are representative of three (A) or two (B, C, H) independent experiments, respectively. Values compared in (A, C-D) using an unpaired Student’s t test (*=p<0.05, **=p<0.005, ***=p<0.0005, ****=p<0.00005).

### B-1a cells differentiate into CD5-IgM secreting cells after stimulation with multiple TLR agonists

Endosomal TLR agonists Imiquimod (TLR7) and ODN CpG 7909 (TLR9) also induced CD5 downregulation on B-1a cells after 3 days of culture (Fig. 3A) as did stimulation with lipids from *Mycobacterium tuberculosis* (Mtb lipids) (Fig. 3C). Similar to LPS stimulation, CD5 expression decreased as the cells divided (Fig. 3B). In contrast, stimulation of CD5+ B-1 cells isolated from mice lacking all TLR-signaling due to a deletion in Unc93, TLR2 and TLR4 (kind gift of Greg Barton, UC Berkeley), were unable to respond with proliferation (Fig. 3D), and failed to loose CD5 (not shown). Thus, B-1 cells were stimulated via TLR-engagement and not via the BCR. In all instances, the loss of CD5 was correlated with the differentiation of CD5+ B-1 cells to IgM-secreting cells, as stimulation of these cells with Imiquimod, CpG 797 and LPS for 3 days resulted in increased percentages of CD138+ cells (Fig. 3E-F) and an increase in IgM concentrations in the culture supernatants (Fig. 3G).

Finally, we examined whether phosphatidylcholine (PTC)-binding B-1 cells can lose CD5 surface expression after TLR-stimulation. PTC is a well-known specificity of a large subset of peritoneal cavity CD5+ B-1 cells ^21, 41^. PTC-binding B-1 cells, identified by incubation of cells with a fluorescent PTC-liposome (kind gift of A. Kantor, Stanford University), lost CD5 expression similarly to PTC non-binders (Fig. 3H). We conclude that TLR-mediated stimulation of CD5+ B-1 cells (“B-1a”) causes the loss of CD5 surface expression, making these cells phenotypically indistinguishable from the proposed B-1 cell “sister” population, the CD5-“B-1b” cells.

### CD5+ B-1 cells become CD5-IgM ASC in the MedLN after Influenza infection

These results raised the possibility that pleural cavity CD5+ B-1 cells respond to influenza virus infection with proliferation, down-regulation of CD5 and differentiating into IgM-secreting cells in the MedLN, explaining the increases in CD5-B-1 cells in the MedLN after influenza infection (Fig. 1), despite their inability to enter these lymph nodes ^33, 34^.

To test this hypothesis, neonatal B-1 cell chimeras generated with FACS-purified CD5+ and CD5-B-1 cells (Igh^b^) at different ratios were infected with influenza virus for 7 days and analyzed for the presence of CD5+ and CD5-MedLN B-1 cells and their ability to secrete IgM (Fig. 4A). As shown previously ^33^, MedLN of mice reconstituted with CD5-B-1 cells had reduced MedLN B-1 cell after infection (Fig. 4B) The frequency of CD5+ and CD5-cells among total donor B-1 cells in the MedLN was similar, regardless of the initial percentage of CD5+ cells (Fig. 4C-D). Of particular note, in chimeras generated with only CD5+ B-1, >50% of B-1 cells in the MedLN lacked CD5 expression (Fig. 4D).

**Figure 4:**
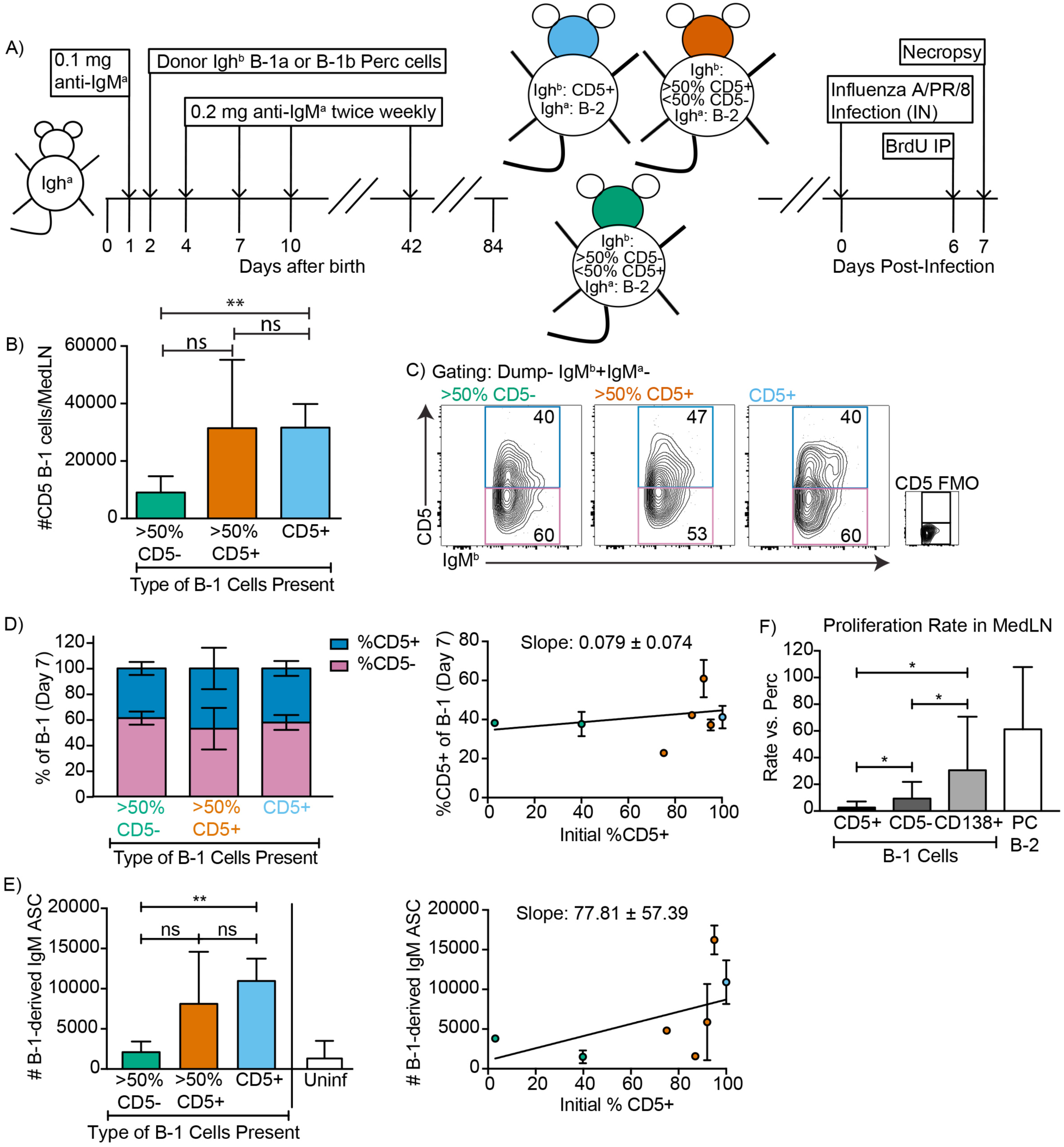
CD5+ B-1 cells differentiate to CD5-IgM ASC in the MedLN after Influenza infection. (A) Neonatal chimeric mice were generated with FACS sorted CD19+ CD23-Ighb+ CD5+ (100%, blue), mostly CD5+ (orange), or mostly CD5- (green) peritoneal cavity-derived B-1 cells and infected with influenza A/Puerto Rico 8/34 for 7 days. (B) Mean number ± SD of B-1 cells in the MedLN of mice 7 days after infection. (C) FACS plot and (D) mean percentage ± SD of Dump-IgMb+ IgMa-CD5+ and CD5-MedLN B-1 cells on day 7. CD5 FMO (fluorescence minus one) control for CD5. (D) Mice were grouped by initial percentage of CD5+ and CD5-B-1 cells (left) and % MedLN CD5+ B-1 cells present on days 0 (initial %) and 7 of infection were plotted with a line of best fit (right). (E) Mean B-1 derived IgM ASC ± SD per MedLN, grouped by initial percentage of CD5+ and CD5-cells (left) and plotted based on initial starting percentage of CD5+ cells (right) with a line of best fit. (F) Mean proliferation rate per day ± SD of CD5+, CD5-, and CD138+ B-1 cells and CD138+ B-2 cells (B-2 PC) in the MedLN compared to proliferation rate per day of similar populations (B-1 or B-2 cells) in the peritoneal cavity of each mouse as determined by BrDU incorporation. Results for infected mice in (B-F) are combined from 4 independent experiments (n=4 for >50% CD5-, n= 7 for >50% CD5+ cells, n=5 for pure CD5+ cells). Results for uninfected chimeras in (E) are combined from 3 independent experiments, n=6. Values in (B, D-F) were compared by unpaired Student’s t test (*=p<0.05, ***=p<0.0005).

Allotype-specific ELISPOTs showed that chimeric mice generated with only CD5+ B-1 cells formed significantly higher frequencies of B-1-derived IgM-secreting cells compared to chimeric mice generated with predominantly CD5-B-1 cells (Fig. 4E). In fact, chimeras generated with CD5-B-1 cells showed no more B-1 derived IgM ASC in their MedLN than uninfected chimeras, consistent with their deficiency in entering the MedLN after infection (Fig. 4E, left panel). There was a significant positive correlation between the frequencies of CD5+ B-1 cells transferred to generate the neonatal chimeras and the ability of the mice to generate IgM ASC following influenza virus infection (Fig. 4E, right panel).

CD5+ B-1 cells failed to show signs of clonal expansion following their accumulation in the MedLN ^33^, which was confirmed using BrdU injection on day 6 after infection. However, the CD5-MedLN B-1 cells showed increased proliferation compared to their counterparts in body cavities (Fig. 4F). Among B-1 cells the proliferation rate was highest among the CD138+ B-1PC, with rates similar to that of the B-2 CD138+ plasma cell compartment (Fig. 4F). The data support the hypothesis that CD5-B-1 and B-1PC arise from proliferating CD5+ pleural cavity B-1 cells that accumulate in the MedLN and differentiate into CD5-IgM ASC following influenza virus infection.

### CD5+ B-1 cells become CD5-IgM ASC in the Mesenteric LNs and Peyer’s Patches after Salmonella typhimurium infection

Numerous infection models have reported CD5-“B-1b” cell responses after infection, including studies on mice infected with *Streptococcus pneumonia* ^29^ and *Salmonella typhimurium* ^32^. This has led to the concept that the CD5-B-1b are a “responder” B-1 cell population, whereas CD5+ B-1 cells generate natural IgM exclusively in the steady state ^29, 42^. We therefore aimed to reexamine whether activation and differentiation of CD5+ B-1 cells into CD5-IgM ASC were more universal outcomes of CD5+ B-1 cell activation to infections.

Neonatal chimeric mice generated with varying ratios of FACS-purified CD5+ and/or CD5-B-1 cells, as described above were orally infected with *Salmonella typhimurium* (Fig. 5A). Again we found a similar percentage of CD5+ and CD5-B-1 cells (identified as IgM^b^+IgM^a^-) in the Mesenteric LN and Peyer’s Patches on day 4 after infection, regardless of the initial percentage of CD5+ cells used to reconstitute the B-1 compartment of these mice (Fig. 5B-C). Over 50% of the B-1 cells in tissues of chimeras established with only CD5+ B-1 cells lacked CD5 surface expression (Fig. 5B-C) and these chimeras were the most competent at forming IgM secreting cells after infection (Fig. 5D-E). In contrast, the MesLN and PP of chimeric mice that were given primarily B-1b cells formed fewer IgM secreting cells, although more than the uninfected chimeras (Fig. 5D-E).

**Figure 5:**
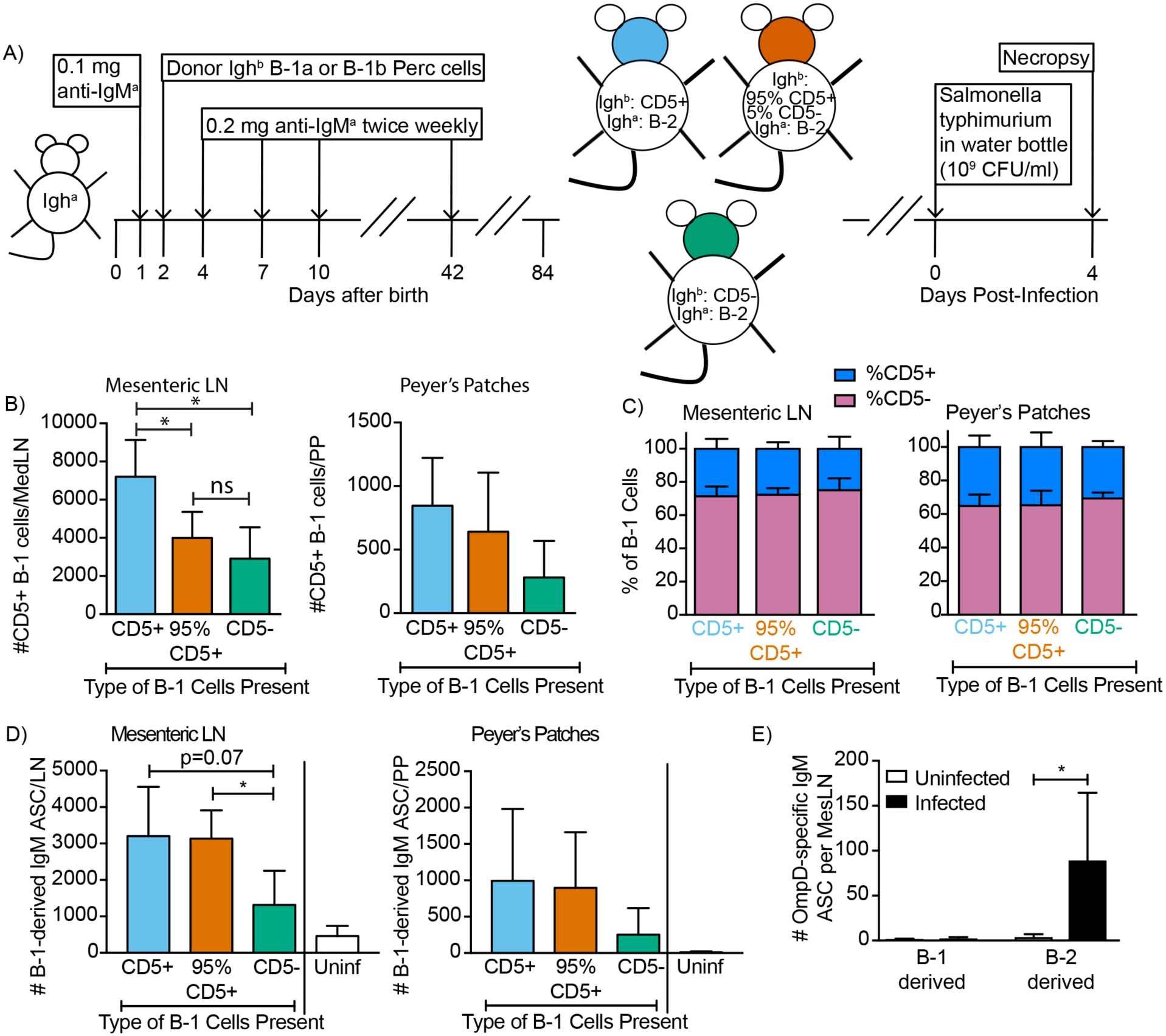
CD5+ B-1 cells differentiate to CD5-IgM ASC in the MesLN, Peyer’s patches, and spleen after *S*. *typhimurium* infection. (A) Neonatal chimeric mice were generated with FACS sorted Dump-CD19+ CD23-Ighb+ CD5+ (100%, blue), CD5- (98%, green), or mostly CD5+ (orange) peritoneal cavity B-1 cells. Mean percentage ± SD of CD5+ and CD5-B-1 cells (IgMb+IgMa-) in (B) MesLN and (C) Peyer’s Patches (PP) on day 4 after oral infection with Salmonella typhimurium via drinking water (n= 3, CD5-; n=4, 95% CD5+; n=6, CD5+). Mean B-1 derived IgM ASC ± SD per (D) MesLN and (E) PP (n= 3, CD5-; n=4, 95% CD5+; n=6, CD5+, uninfected). (F) B-1 and B-2 derived OmpD-binding IgM ASC per MesLN in uninfected and infected neonatal chimeric mice (n=5-6). Results in (B-F) are combined from 2 independent experiments, uninfected chimeras in (D) are combined from 3 independent experiments. Values in (B-F) were compared with an unpaired Student’s t test (*=p<0.05)

The *S*. *typhimurium* surface antigen OmpD had been reported previously to stimulate IgM secretion exclusively by CD5-“B-1b” cells ^32^. However, in our hands, although total B-1-derived IgM was increased after infection in the Mesenteric LN (Fig. 5D), OmpD-specific B-1-derived IgM ASC did not increase significantly (Fig. 5F). Instead, we found OmpD-specific IgM secretion by B-2 derived plasmablasts. Of note, the phenotype of B-2-derived plasmablasts is indistinguishable from that of “B-1b” cells (CD19low CD45lo IgM+ CD43+ (Fig. 5F)), and thus only identifiable using a lineage-marking approach, such as used here.

Together these findings demonstrate that in response to both bacterial and viral infections, B-1 cells accumulating in secondary lymphoid tissues lose CD5 expression and become the main source of B-1 derived IgM secretion after both bacterial and viral infections. *In vitro* this process was recapitulated by stimulation with various TLR-ligands.

### Changes in BCR signaling following innate activation of B-1 cells

Surface CD5 expression by B-1 cells has been linked previously to their inability to proliferate in response to BCR-mediated signaling ^25^. To analyze the association of CD5 with the BCR on B-1 cells in steady state and to determine what alterations stimulation of the BCR may induce on B-1 cells, we analyzed the IgM-BCR-complexes on the cell surface of highly FACS-purified, then rested, peritoneal cavity CD5+ CD45R^lo^ CD23-B-1 and splenic CD45R^hi^ CD23+ CD5-follicular B cells using Proximal Ligation Assay (PLA). Both CD19 and CD5, were strongly associated with the surface-expressed IgM-BCR on B-1 cells, while CD5 was not directly associated with the co-stimulator and signaling molecule CD19 (Fig. 6A). Consistent with the lack of stimulation and strong interaction between IgM and CD5, the BCR-signaling chain CD79 only weakly interacted with the adaptor molecule Syk in B-1 cells in the steady-state. B-2 cells lack CD5 expression, and CD19 did not interact with the IgM-BCR prior to stimulation (Fig. 6A).

**Figure 6:**
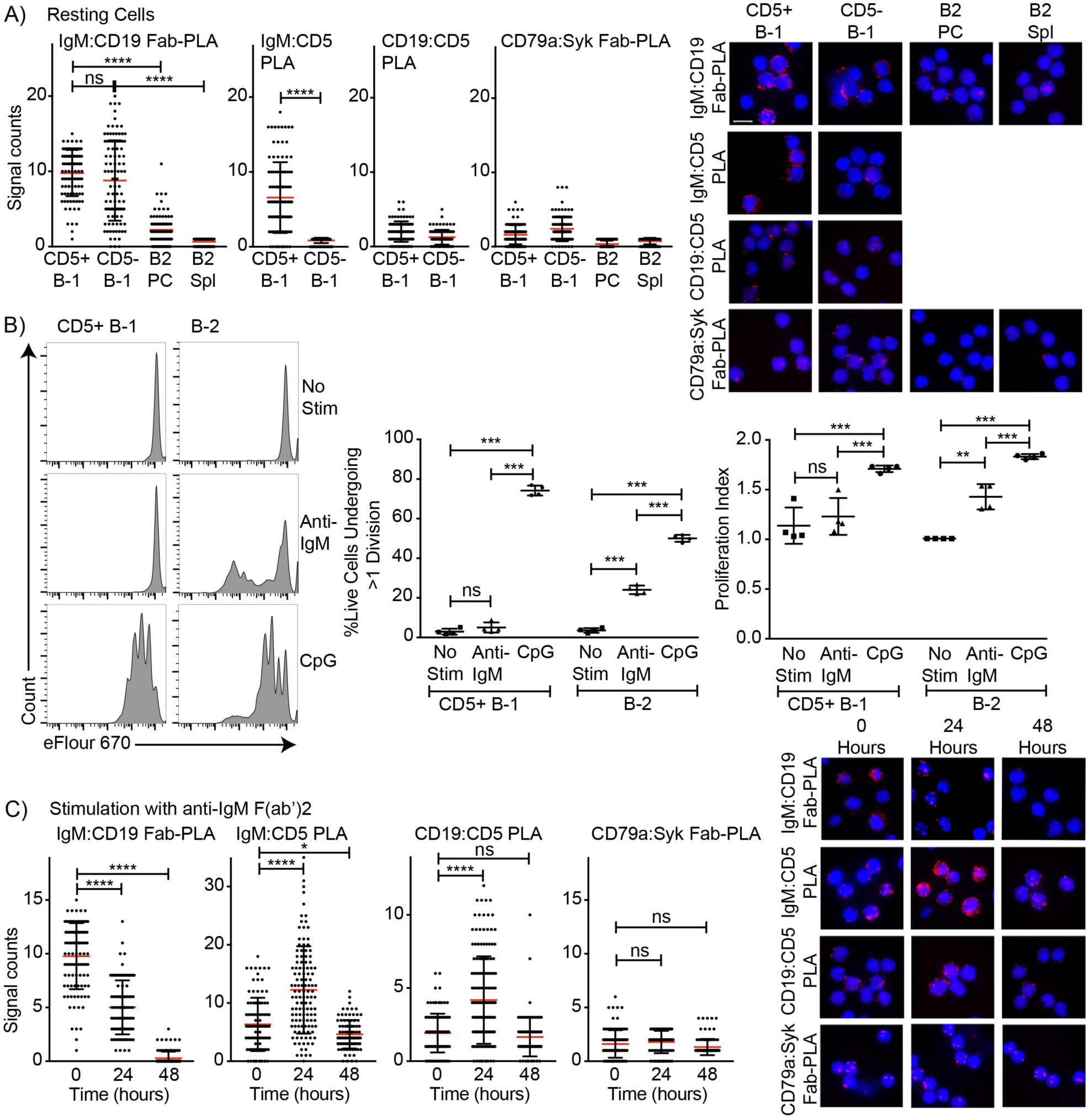
Association of CD5 with IgM-BCR in resting B-1a cells is increased after BCR-stimulation. **(A)** Indicated FACS-purified B cell subsets from the peritoneal cavity (PC) and spleen (Spl) of C57BL/6 mice were analyzed by proximal ligation assay for the following interactions (left to right): IgM:CD19, IgM:CD5, CD19:CD5 and CD79:syk. Left panel shows summarizes data on signal counts for 200 individual cells analyzed. Each symbol represents one cell, horizontal line indicates mean signal count per cell. Right panel show representative fluorescent images. (**B**) FACS-purified CD19hi CD23-CD5+ CD43+ B-1 cells from the peritoneal cavity and CD19+ CD23+ splenic B-2 cells were labeled with efluor670 and then cultured in the absence (top) or presence of 20 ug/ml anti-IgM (middle) or 10 ug/ml CpGs for 72h. Left panels show representative histogram plots, middle panel shows the % cells in each culture having undergone at least one cell division and right panel indicates the proliferation index (average number of proliferations undergone per divided cell). (**C)** Summary of proximal ligation assay results of B-1 cells purified as in (A) and then stimulated for indicated times with anti-IgM(Fab)2. Interactions of the following proteins were analyzed on 200 cells per condition (left to right): IgM:CD19, IgM:CD5, CD19:CD5 and CD79:syk. Right panels shows representative fluorescent images from one experiment of at least two done. Values were compared using an unpaired Student’s t test (*=p<0.05, **=p<0.005, ***=p<0.0005, ****=p<0.00005).

As expected, stimulation of CD5+ B-1 cells with anti-IgM failed to induce their proliferation but did induce proliferation of B-2 cells. In contrast, stimulation with CpG (TLR9-ligands) induced strong proliferation by both B-1 and B-2 cells (Fig. 6B). The lack of responsiveness to BCR-stimulation was explained by the PLA data, which showed the maintenance and even increase in IgM-BCR:CD5 association and an increase in the association of the inhibitor CD5 with CD19 following anti-IgM treatment. Furthermore, B-1 cells lost the interaction of the IgM-BCR with CD19. Consequently, CD79-Syk interaction remained very low (Fig. 6C). Thus, BCR-engagement on B-1 cells inhibits BCR-signaling by reducing steady-state IgM-CD19 interactions and likely also by initiating instead interactions between CD5 and CD19.

In contrast to direct stimulation of the IgM-BCR, CpG stimulation led to changes in the BCR-signaling complex that are consistent with induction of positive BCR-signaling, and/or the ability to signal through the antigen-receptor. CpG stimulation strongly increased the interaction of IgM-BCR:CD19 and reduced CD5:IgM-BCR proximity. The already weak CD5:CD19 interaction was further reduced (Fig. 7A), consistent with the loss in surface CD5 expression noted following stimulation (Fig 3). These rapid changes in the BCR-signaling complex were associated with increases in CD79:Syk interaction, suggesting active BCR downstream signaling in CpG-stimulated B-1 cells. This was further supported by sustained increased levels of phosphorylated Akt (pAkt; pS473), while anti-IgM stimulation reduced pAkt levels below that of unstimulated B-1 cells by 24h, after an initial increase (Fig. 7B).

**Figure 7:**
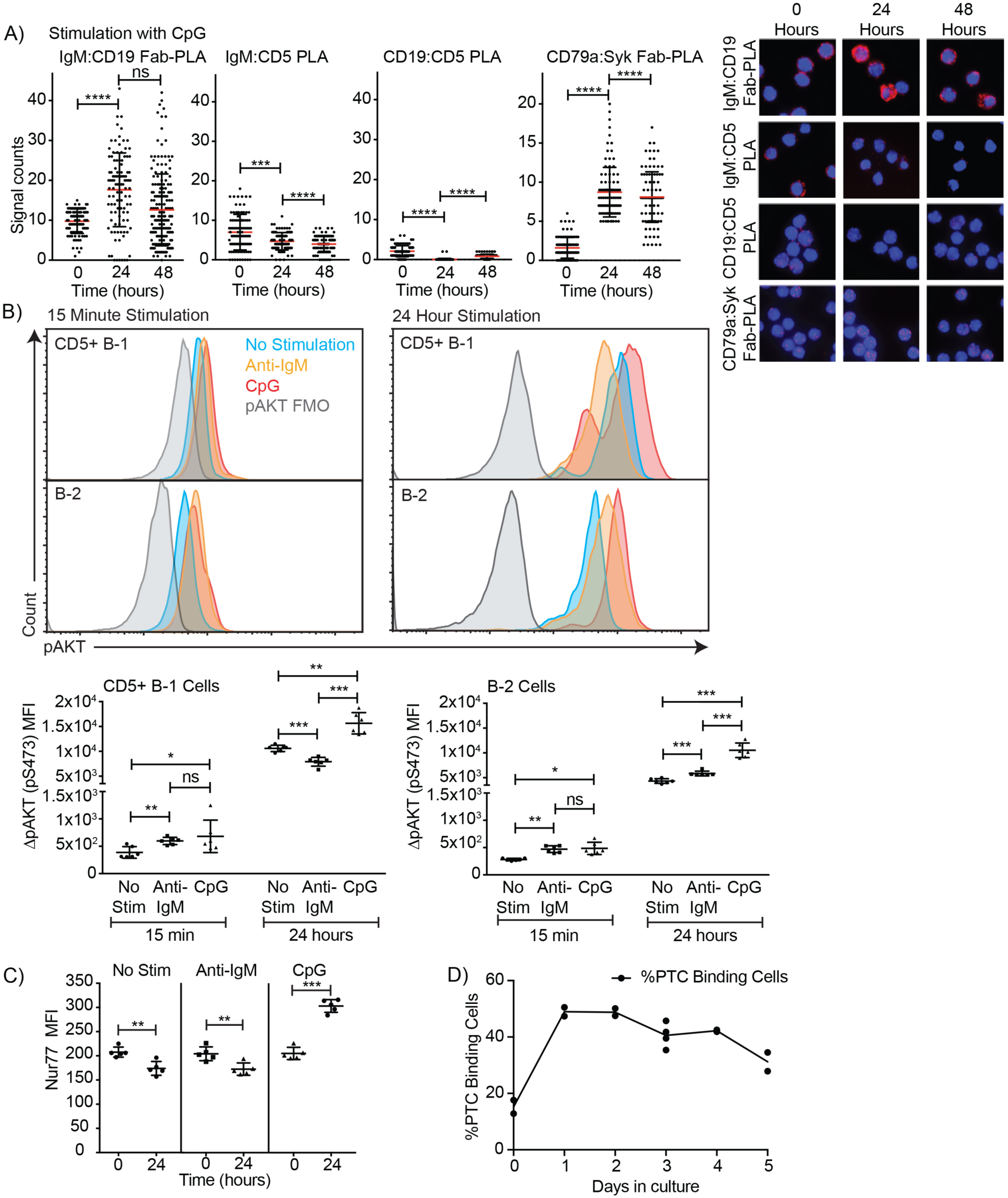
TLR-mediated stimulation of CD5+ B-1 cells alters the BCR-signalosome. **(A)** FACS-purified peritoneal cavity CD19^hi^ CD23-CD43+ CD5+ B-1 and splenic CD19+ CD23+ CD43-CD5-B-2 cell of C57BL/6 mice were stimulated for the indicated times with TLR9-agonist ODN7909 prior to analysis by proximal ligation assay, probing for the following interactions (left to right): IgM:CD19, IgM:CD5, CD19:CD5 and CD79:syk. Left panel summarizes data on signal counts for 200 individual cells analyzed. Each symbol represents one cell, horizontal line indicates mean signal count per cell. Right panel show representative fluorescent images. (**B**) Analysis of the phosphorylation status of Akt by probing for Akt pS473 by flow cytometry. Top panels show representative histogram plots, bottom summarizes the results. (**C**) Mean fluorescence intensity ± SD of staining for the immediate early activation factor Nur77, in CD5+ B-1 cells isolated as described in (A) and cultured for up to 2 days in the absence and presence of the indicated stimuli. (**D**) Shown are % frequencies of live PtC-binding B-1 cells among live FACS-purified CD5+ peritoneal cavity B-1 cells cultured with LPS stimulation for the indicated times, as assessed by flow cytometry. Each symbol represents results obtained from one culture well. Results are representative from experiments conducted at least twice with multiple repeats done per experiment (n=2-5). Results in D are combined from two independent experiments. Values were compared using an unpaired Student’s t test (*=p<0.05, **=p<0.005, ***=p<0.0005).

We also noted increased Nur77 expression in CpG-but not anti-IgM-stimulated B-1 cells at 24h (Fig. 7C) but not at 3h (not shown). Given that B-1 cell responses following CpG-stimulation were TLR-expression dependent *in vitro* (Fig. 3D), and no external antigen was provided to the cultures, TLR-signaling may directly link to BCR-signaling, or TLR-signaling may “license” subsequent self-antigen recognition, by altering the BCR-signaling complex. In support of the latter, we noted a strong increase in the frequencies of PTC-binding B-1 cells during culture (Fig. 7D), which may be due to the expansion of CpG-activated B-1 cell in response to PTC-antigen present on dead and dying cells in the cultures.

### Local IgM production following influenza infection depends on TLR expression

TLR-expression was also required for B-1 cell responses to influenza virus infection in vivo, as complete TLR-/- mice (due to a lack of TLR2, TLR4 and Unc93) showed significant deficits in CD5+ B-1 cell responses following infection (Fig. 8A/B), resulting in a significant increase in the ratio of CD5+ over CD5- and CD19+ CD43+ B cells. This suggested that CD5+ B-1 cells in TLR-/- mice could enter the MedLN, consistent with our previous findings that this step is TLR-independent but Type I IFN-dependent ^34^, but once in the MedLN were not activated via TLR-dependent signals to downregulate CD5 (Fig. 8B). Of importance, the lack of TLR-stimulation also resulted in a near complete loss of CD19^lo/-^ IgM+ CD138+ B-1PC in the MedLN at day 5 of infection (Fig. 8C), and a corresponding drop in IgM ASC in TLR-/- compared to control mice at that timepoint (Fig. 8D), while viral loads were similarly low in the lungs of both mouse strains (not shown). Generation of Ig-allotype chimeras in which only B-1 cells lacked TLR expression confirmed a B-1 cell-intrinsic requirement for TLR-signaling in B-1 cell differentiation to CD138+ ASC after influenza virus infection (Fig. 8D-F).

**Figure 8:**
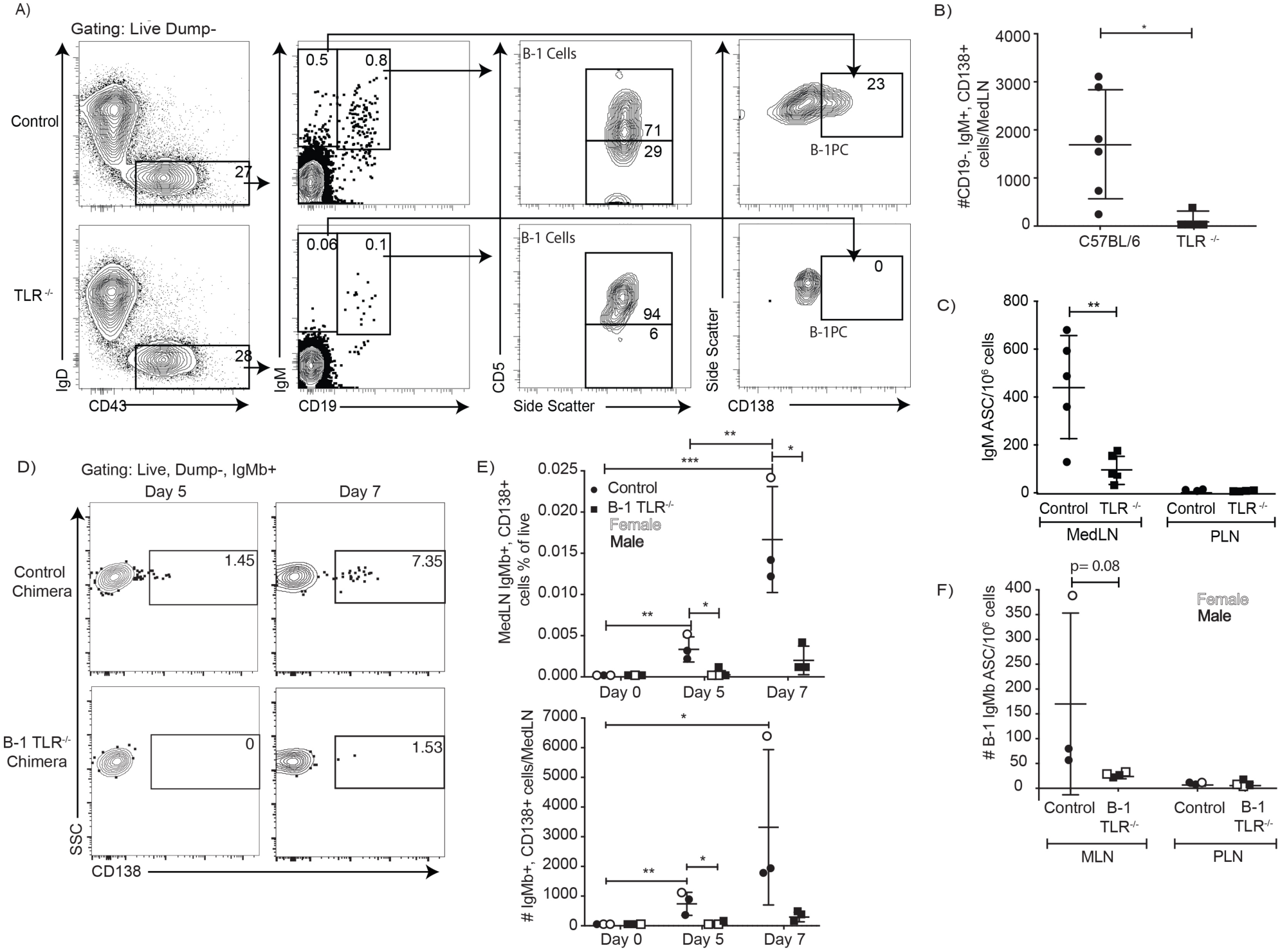
TLR-mediated stimulation is required for maximal IgM responses to influenza virus infection. (**A**) C57BL/6 (n = 5) and congenic total TLR-/- mice (n =5; lacking TLR2, TLR4 and Unc93) were infected with influenza A/Puerto Rico/8/34 for 5 days. Shown are representative FACS plots from control C57BL/6 (top) and TLR-/- (bottom) mice FACS analysis of MedLN for the presence of B-1 and B-1PC. (**B**) Total number of CD19-IgM+ CD138+ plasmablasts as assessed by FACS and (**C**) IgM-secreting cells per MedLN (or PLN as controls) as assessed by ELISPOT of infected mice. (**D-F**) Similar analysis as for A-C but using allotype chimeras generated with wild type recipients and B-1 cells from either C57BL/6 or TLR-/- mice. (D) Representative FACS analysis of CD138+ B-1PC pre-gated for live, dump-, B-1 donor (IgMb+) cells in MedLN on days 5 and 7 after influenza infection. (**E**) Mean ± SD of data summarized from analysis shown in D. (F) Mean ± SD of B-1 IgM-ASC in MedLN on day 7 after infection, as assessed by ELISPOT. Each symbol represents results from one mouse with female mice shown as open symbols, males as closed symbols. Results are combined from two independent experiments. Values were compared using an unpaired Student’s t test (*=p<0.05, **=p<0.005, n.s. not significant).

Together the data demonstrate a linkage of TLR and BCR-signaling during B-1 cell responses to infections, with intrinsic TLR-mediated signaling triggering a rapid reorganization of the IgM-BCR-signalosome complex including the removal of the BCR-signaling inhibitor CD5 and the increased association of IgM-BCR with the co-receptor CD19, resulting in the differentiation of CD5+ B-1 to CD5-IgM-secreting B-1 and B-1PC.

## Discussion

Self-reactive, fetal and neonatal-developing B-1 cells are not activated readily in response to BCR-stimulation, owing to an unresponsive IgM-BCR complex, previously shown to be due to expression of the BCR-signaling inhibitor CD5 and a lack of fully functional CD19 signaling. Yet B-1 cells respond rapidly to various infections with migration to secondary lymphoid tissues and differentiation into IgM secreting cells. The present study suggests a mechanism by which B-1 cell can overcome their inherent BCR-signaling block, namely via TLR-mediated reorganization of the BCR-signalosome complex. This non-redundant signal was shown to cause the loss of CD5 association with the IgM-BCR and eventually CD5 expression itself, the increased association between IgM and the co-stimulatory molecule CD19, and a resulting strong increase in CD79:Syk interaction and phosphorylation of BCR downstream effectors.

Thus, TLR-signaling might not simply only activate B-1 cells innately, but may also activate or “license” B-1 cells for subsequent BCR-mediated signaling. This could explain the recent data by Kreuk and colleagues, which associated specific TLR-signaling of B-1 cells with antigen-specific B-1 cell responses. Most notable was the early loss of CD5:BCR association and eventual loss of expression of CD5 on TLR-activated CD5+ “B-1a” cells *in vitro*, and *in vivo* in response to both bacterial (*S*. *thyphimurium*) and viral (influenza virus) infections. The exclusion/removal of the signaling inhibitor CD5 correlated with enhanced CD79:syk interaction, enhanced pAkt levels and the differentiation of B-1 cells. In contrast, IgM-BCR stimulation of CD5+ B-1 cells alone enhanced CD5:BCR interactions and reduced BCR:CD19 interactions, resulting in a failure to induce CD79:syk assocation and sustained phosphorylation of Akt. We conclude that BCR-downstream signaling pathway activation is disabled in self-reactive, CD5+ B-1 cells and that this can be overcome with TLR-mediated activation.

As we showed that it is the CD5+ B-1 cell population that initially responded to infections with influenza virus and with *S*. *typhimurium*, the latter previously identified as an exclusive “B-1b” response ^32, 43^, it appears likely that other pathogen-induced B-1 cell responses may also represent responses of CD5+ B-1 cells ^29, 30, 31, 32, 43^ which subsequently lose CD5, rather than de novo responses of CD5-“B-1b” cells. If true, the lack of CD5-expression on B-1 cells would mark previously activated and differentiated B-1 cells. This is consistent with previous reports on the phenotype of natural IgM-secreting cells ^35, 44^ and could explain also earlier reports that CD5+, not CD5-, B-1 cells form natural IgM secreting cells ^29, 45, 46^. In addition, the data are consistent with findings that CD5-B-1 cells contain CD5-memory-like B-1 cells in the body cavities of previously infected mice ^31, 47^.

The data presented here are inconsistent with models that regard B-1 cell responses as a “division of labor” between two subsets of B-1 cells: B-1a and B-1b cells, where CD5+ B-1a contribute “natural” IgM and CD5-B-1b the induced IgM, proposed previously ^29, 42^. Instead, we show that the CD5-B-1 cells secreting IgM in response to influenza or *Salmonella spp*. were both derived from TLR-activated CD5+ B-1 cells. Since no other clearly subset-defining differences between B-1a and B-1b cells have been reported to-date, it may be more prudent to describe B-1 cells simply as CD5+ and CD5- and to drop the use of the terms B-1a and B-1b, unless and until further clear subset-defining differences are identified.

Our data do not exclude the possibility that some B-1 cells develop which either express low, or undetectable levels of surface CD5, as described previously ^48^. Given the known functions of CD5 as an inhibitor of BCR-signaling ^2, 3, 4, 12^ and the fact that CD5-expression levels on thymocytes correlated with the strengths of the positively selecting TCR–MHC-ligand interactions 4, such CD5lo/neg B-1 cells might have lower levels of self-reactivity ^19, 20, 21, 22^ and lack the need for CD5-mediated silencing of BCR-signaling in order to avoid inappropriate hyperactivation of these self-reactive B cells. De novo development of both CD5+ and CD5-B-1 cells has been reported to occur also in stromal cell cultures seeded with B-1 cell precursors ^49^. It remains to be explored whether the presence of DAMPS in those in vitro cultures could contribute to the loss of CD5 on initially CD5+ B-1 cells, or whether these cells never expressed CD5.

Our data are not consistent with early reports suggesting that CD5+ B-1 cells could only reconstitute themselves, but not CD5-B-1 cells ^48, 50^, as reconstitution of neonatal mice with even very highly FACS-purified body cavity CD5+ B-1 cells led to significant numbers of CD5-B-1 cells recovered from these mice and B-1b (^33^ and Figs. 4/5). This discrepancy remains to be further explored.

The reorganization of the IgM-BCR complex, including the loss of CD5, may allow B-1 cells to gain responsiveness to BCR engagement and subsequent antigen-specific proliferating and differentiation. The activating antigen may be either a foreign antigen, or self-antigens induced at sites of infection and inflammation. The differentiation of B-1 cells during an infection might thus involve a two-step process: A first TLR-mediated signal that renders B-1 cells receptive to BCR signaling and a second antigen-specific signal through antigen engagement with the BCR. Additional positive or negative signals might also be involved.

An alternative model would involve the direct linking of TLR and BCR-signalosome effects. For example, low-affinity BCR-antigen interactions might be supported by having DAMPS and PAMPS first bind to TLR on the B cell surface, which then brings these antigens in close proximity to the BCR, triggering antigen-specific BCR activation event. Or, internalization of antigen-BCR complexes could engage endosomal TLRs. Although we did not provide antigens other than TLR-ligands to our in vitro cultures, dead and dying cells may provide ample DAMPS, including PTC, that could have stimulated these cells. This could explain why we found such strong enrichment for PTC-binders among cultures of TLR-stimulated B-1 cells and the CpG-stimulation dependent activation of the BCR signaling pathways.

The lack of phenotypic differences between CD5-B-1 cells and B-2 cell-derived non-switched plasmablasts, both expressing CD19, CD43 low levels of CD45 and lacking IgD, further complicates the identification of responding B cell subsets *in vivo*, as demonstrated with the analysis to the *Salmonella* antigen OmpD. Using allotype-marked B cell lineages, we show here that anti-OmpD secreting B cells were derived predominantly from B-2 cells in our system, and not as previously suggested from B-1b cells ^32^. When CD5+ B-1 cells lose CD5, they also upregulate CD138, thus becoming indistinguishable from B-2-derived plasma cells and plasmablasts that carry the same phenotype. Some have used expression of CD11b to identify B-1 versus B-2 cells in infectious models ^29, 32^, but this marker is also dynamically regulated depending on tissue site and B-1 cell activation status ^34, 44, 51^. Recent lineage-tracing approaches ^13, 14, 52^ may provide novel approaches and markers that unequivocally identify B-1 cells. In the meantime, the use of neonatal B-1 allotype-chimeras remains a valuable tool for such analyzes.

Taken together, our data suggest that the BCR-complex composition on neonatally-derived, self-reactive B-1 cells is controlled by TLR-mediated signals, preventing inappropriate activation and autoimmune disease on the one hand, while facilitating rapid B-1 cell participation in anti-viral and anti-bacterial infections on the other. TLR-signaling thereby influences not only innate but also antigen-specific B-1 cell activation.

## Materials and Methods

### Mice

8-16 week old male and female C57BL/6J mice and breeding pairs of B6.Cg-Gpi1^a^Thy1^a^Igh^a^/J (Igh^a^) were purchased from The Jackson Laboratory. Female, 10 weeks old BALB/C mice were purchased from Jackson Laboratory. B6-Cg-Tg(PRDM1-EYFP)^1Mnz^ (Blimp-1 YFP) breeders were kindly provided by Michel Nussenzweig (The Rockefeller University, NY) and breeding pairs of Tlr2^-/-^ x Tlr4^-/-^ xUnc93b1^3d/3d^ (TLR-/-) mice by Greg Barton (University of California, Berkeley, CA). Mice were housed under SPF conditions in micro-isolator cages with food and water provided ad libitum. Mice were euthanized by overexposure to carbon dioxide. All procedures were approved by the UC Davis Animal Care and Use Committee.

### Chimera generation

Neonatal chimeric mice were generated as described previously ^53^. Briefly, one-day old Igha C57BL/6 congenic mice were injected intraperitoneally with anti-IgM^a^ (DS-1.1) diluted in PBS. On day 2 or 3 after birth mice were injected with total peritoneal cavity wash out, or with FACS-purified dump-CD19+ CD43+ CD5+ and/or CD5-B-1 cells from C57BL/6 or TLR-/- mice (Igh^b^) mice. Intraperitoneal anti-IgM^a^ injections were continued twice weekly until mice reached 6 weeks of age. Mice were then rested for at least 6 weeks before use, for reconstitution of the conventional B cell populations from the host bone marrow.

### Influenza virus infection

Influenza A/Puerto Rico/8/34 was grown and harvested as previously described ^54^. Mice were anesthetized with isoflurane and virus was diluted to a previously titrated sublethal dose of infection and administered intranasally in PBS.

### Salmonella typhimurium infection

Oral infections with *S*. *typhimurium* were performed following previously described protocols ^55^. *S*. *typhimurium*, strain SL1344, kindly provided by Stephen McSorley (University of California, Davis, CA), was grown overnight at 37°C in Luria-Bertani broth. A known volume of bacteria were centrifuged for 20 minutes at 6,000-8,000 rcf at 4°C after concentration was determined by spectrophotometer reading at OD_600_. Bacterial pellets were resuspended in mouse drinking water to a concentration of 10^9^ CFU/ml. Water was provided to mice ad lib.

### Flow cytometry and sorting

Tissues were processed and stained as described previously ^56^. Briefly, single cell suspensions of spleen, lymph node, and Peyer’s patches were obtained by grinding tissues between the frosted ends of two microscope slides, then resuspended in “Staining Media” ^56^. Peritoneal cavity washout was obtained by introducing Staining Media into the peritoneal cavity with a glass pipet and bulb, agitating the abdomen, and then removing the media. Samples were filtered through nylon mesh and treated with ACK lysis buffer as needed. Cell counts were performed using Trypan Blue exclusion to identify live cells.

Fc receptors were blocked using anti-CD16/32 antibody (2.4G2) and cells were stained using fluorochrome conjugates generated in-house unless otherwise specified against the following antigens: CD19 (clone ID3)-Cy5PE, allophycocyanin, FITC, CD4- (GK1.5), CD8a- (53- 6.7), CD90.2- (30H12.1), Gr1- (RB68-C5), F4/80- (F4/80), and NK1.1- (PK136) Pacific blue (“Dump”), CD43- (S7) allophycocyanin or PE, IgM- (331) allophycocyanin, Cy7-allophycocyanin, FITC, Alexa700, IgM^a^- (DS-1.1) allophycocyanin, biotin, IgM^b^- (AF6-78.2.5) allophycocyanin, FITC, biotin, CD5- (53-7.8) PE, biotin, IgD- (11-26) Cy7PE, Cy5.5PE, CD138- (281-2) allophycocyanin, PE; CD138-BV605 (BD Bioscience), CD19-BV786, PE-CF594 (BD Bioscience), SA-Qdot605 (Invitrogen), SA- allophycocyanin (eBioscience), BrDU-FITC (BD Bioscience), B220 (CD45R) APC-eFluor 780 (Invitrogen) and CD23-biotin (eBioscience), BV605, BV711 (BD Bioscience). PTC-FITC liposomes were a kind gift of Aaron Kantor (Stanford University, CA). Dead cells were identified using Live/Dead Fixable Aqua or Live/Dead Fixable Violet stain (Invitrogen).

Intracellular staining: Cells were surfaced stained, then fixed (eBiosience IC Fixation Buffer) for 30 minutes and then permeabilized (eBioscience Permeablization Buffer) for 30 minutes, followed by staining with anti-Nur77-Alexa Fluor 488 for 30 minutes all at room temperature.

Phosphoflow: Cells were fixed (BD Cytofix) for 12 minutes at 37°C. Cells were then permeabilized (BD Perm Buffer III) for 30 minutes on ice and intracellularly stained with anti-phospho-Akt-Alexa Fluor 488 for 30 min on ice.

FACS analysis was done using either a 4-laser, 22-parameter LSR Fortessa (BD Bioscience) or a 3-laser FACSAria (BD Bioscience). Cells were sorted as previously described ^56^ using the FACSAria and a 100 µm nozzle. Data were analyzed using FlowJo software (FlowJo LLC, kind gift of Adam Treister).

### ELISA

Sandwich ELISA was performed as previously described ^56^. Briefly, MaxiSorp 96 well plates (ThermoFisher) were coated with anti-IgM (Southern Biotech) and nonspecific binding was blocked with 1% NCS/0.1% dried milk powder, 0.05% Tween20 in PBS (“ELISA Blocking Buffer”). Two-fold serial dilutions in PBS of culture supernatants and an IgM standard (Southern Biotech) were added to the plates at previously optimized starting dilutions. Binding was revealed with biotinylated anti-IgM (Southern Biotech), Streptavidin-Horseradish Peroxidase, both diluted in ELISA Blocking Buffer, and 0.005% 3,3’,5,5’-tetramethylbenzidine (TMB)/0.015% hydrogen peroxide in 0.05 M citric acid. The reaction was stopped with 1N sulfuric acid. Antibody concentrations were determined by measuring sample absorbance on a spectrophotometer (SpectraMax M5, Molecular Devices) at 450 nm (595 nm reference wavelength) and then compared to a standard curve created with a mouse IgM standard (Southern Biotech) of known concentration.

### Culture and proliferation dye labeling

After FACS sorting, cells were labeled with Efluor670 or CFSE at previously determined optimal concentrations, by incubation at 37°C for 10 mins., then washed three times with staining medium containing 10% neonatal calf serum and resuspended into “Culture Media” (RPMI 1640 with 10% heat inactivated fetal bovine serum, 292 µg/ml L-glutamine, 100 Units/ml penicillin, 100 µg/ml streptomycin, and 50 µM 2-mercaptoethanol). Cells were plated at 105 cells/well of 96-well U bottom tissue culture plates (BD Bioscience), and unless otherwise indicated, cultured at 37°C/5% CO_2_ for 3 days. When indicated, LPS at 10 µg/ml, Mycobacterium TB lipids at 20 µg/ml (BIA), Imiquimod (R837, InvivoGen) at 1 µg/ml, CpG ODN 7909 at 5 µg/ml or anti-IgM (Fab)_2_ at 10-20ug/ml were added to the wells. Cell enumeration after culture was performed using Molecular Probes CountBright Beads (Thermo Fisher) by flow cytometry, per manufacturer instructions. After culture, culture plates were spun and supernatant was collected and stored at −20°C, and cells were stained for FACS.

### ELISPOT

IgM antibody secreting cells were enumerated as previously described ^54^. Briefly, 96 well ELISPOT plates (Multi-Screen HA Filtration, Millipore) were coated overnight with anti-IgM (331) or recombinant OmpD (MyBioSource) and non-specific binding was blocked with 4% Bovine Serum Albumin (BSA)/PBS. Cell suspensions were processed, counted, and directly plated in culture medium into ELISPOT wells and subsequently serially diluted two-fold, or they were FACS-sorted directly into culture media-containing ELISPOT wells. Cells were incubated overnight at 37°C/5% CO_2_. Binding was revealed with biotinylated anti-IgM (Southern Biotech), anti-IgM^a^ (BD Bioscience), or anti-IgM^b^ (BD Bioscience), Streptavidin-Horseradish Peroxidase (Vector Labs) both diluted in 2% BSA/PBS, and 3.3 mg 3-amino-9-ethylcarbazole (Sigma Aldrich) dissolved in dimethyl formamide/0.015% hydrogen perioxide/0.1M sodium acetate. The reaction was stopped with water. Spots were enumerated using the AID EliSpot Reader System (Autoimmun Diagnostika, Strassberg, Germany).

### qRT-PCR

mRNA was isolated from cells using the RNeasy mini kit (Qiagen), per manufacturer instructions. cDNA was generated using random hexamer primers and SuperSript II reverse transcriptase (Invitrogen). qRT-PCR was performed using commercially available Taqman primer/probes for *cd5* and *ubc* (Thermo Fisher).

### BrDU labeling

Mice were injected intraperitoneally with 1 mg of BrDU (Sigma-Aldrich) per mouse diluted in 100 µL PBS, 24 hours before tissue collection. Staining for BrDU was performed as described previously ^56^.

### Proximity Ligation Assay (PLA)

After FACS sorting, cells were resuspended in RPMI and rested for at least two hours before designated stimuli were added to culture media. Stimulated and unstimulated cells were cultured for 5 minutes, and 24 and 48 hours prior to PLA. PLA was performed as previously described ^57^. In brief: For PLA-probes against specific targets, the following unlabeled Abs were used: anti-IgM (Biolegend, clone RMM-1), anti-CD79a (Thermo Fisher, clone 24C2.5), anti-CD5 (Biolegend, clone 53-7.3), anti-Syk (Biolegend, clone Syk-01), and anti-CD19 (Biolegend, clone 6D5). Fab fragments against CD79a, Syk, IgM, and CD19 were prepared with Pierce Fab Micro preparation kit (Thermo Scientific) using immobilized papain according to the manufacturer’s protocol. After desalting (Zeba spin desalting columns, Thermo Scientific), all antibodies were coupled with PLA Probemaker Plus or Minus oligonucleotides (Sigma-Aldrich) to generate PLA-probes. For in situ PLA, B cells were settled on polytetrafluoroethylene slides (Thermo Fisher Scientific) for 30 min at 37 °C. BCR. Cells were fixed with paraformaldehyde 4%, for 20 min. For intracellular PLA, B cells were permeabilized with 0,5% Saponin for 30 min at room temperature, and blocked for 30min with Blocking buffer (containing 25 μg/ml sonicated salmon sperm DNA, and 250 μg/ml bovine serum albumin). PLA was performed with the Duolink In-Situ-Orange kit. Resulting samples were directly mounted on slides with DAPI Fluoromount-G (SouthernBiotech) to visualize the PLA signals in relationship to the nuclei. Microscope images were acquired with a Leica DMi8 microscope, 63 oil objective (Leica-microsystems). For each experiment a minimum of 100 B-1a/B-1b/B-2 peritoneal cavity or 1000 splenic B-2 cells from several images were analyzed with CellProfiler-3.0.0 (CellProfiler.org). Raw data were exported to Prism7 (GraphPad, La Jolla, CA). For each sample, the mean PLA signal count per cell was calculated from the corresponding images and the statistical significance with Mann–Whitney test.

### Statistical analysis

Statistical analysis was done using a two-tailed Student t test with help of Prism software (GraphPad Software). For time-course data, an ANOVA was performed with the help of Prism software, and if significant, Student t tests were performed to determine which time points were significant. When multiple comparisons were run on the same sets of data, Holm-Sidak correction was applied, using Prism software. p < 0.05 was considered statistically significant.

## Acknowledgements

We thank Drs. Joseph Benoun and Stephen McSorley (UC Davis) for expert help in conducting infections with *S*. *typhimurium*, Dr. Aaron Kantor (Stanford University) for PtC liposomes, and Dr. Greg Barton (UC Berkeley) for TLR-/- mice and discussions. This work was supported through NIH/NIAID grants U19-AI109962 (N.B) and R01-117890 (N.B.), the National Center for Advancing Translational Sciences, NIH, through grant number UL1 TR000002 and linked award TL1 TR000133 (H.P.S), the NIH-2T32OD010931-09 (H.P.S), NIH – 5T35OD010956 (H. P.S) and the T-32 AI060555 (H.P.S) and NIH-T32 OD011147 (F.L.S).

